# Evaluating DNA metabarcoding to characterise diet diversity and foraging strategies in a generalist mesopredator, the lesser black-backed gull (*Larus fuscus*)

**DOI:** 10.64898/2026.07.17.738687

**Authors:** Alice Risely, David N. Carss, Emma F. How, Bart J. Donato, Tim D. Frayling, Elnaz Poulab, Naiara Guimarães Sales

## Abstract

1. Information on diet composition at both individual and population levels is fundamental to understanding resource use and availability, which influence individual fitness and the population dynamics of both predators and their prey. DNA diet metabarcoding offers a powerful molecular approach to diet analysis, but its application can be limited by methodological biases, particularly in generalist species with highly diverse diets.
2. Here, we evaluate the performance of DNA metabarcoding for diet analysis in the lesser black-backed gull (*Larus fuscus*), a highly generalist mesopredator whose populations have shown complex responses to changes in natural and anthropogenic food resources over the past half-century. We collected faecal and regurgitate samples from pre-fledging chicks at two coastal and inland breeding colonies in northwest England, alongside pharyngeal, stomach, intestinal, and cloacal swabs from adult gull carcasses. Samples were analysed using COI (targeting animal DNA) and 12S (targeting vertebrate DNA) metabarcoding markers, and three blocking primers were developed to reduce host amplification in 12S libraries.
3. Metabarcoding performance varied substantially among sample types and primer combinations. Without blocking primers, usable dietary information was recovered from regurgitate and stomach samples but not from intestinal or faecal samples. Blocking primers improved recovery of dietary DNA from faecal samples, but also increased the amplification of contaminants, elevating the risk of false-positive detections.
4. Across all sample types, metabarcoding identified 71 unique species belonging to 61 genera, including earthworms, small mammals, commercial and non-commercial fish species, lapwing, and urban-derived food products originating from livestock species. Dietary profiles revealed distinct clusters of consumed species associated with urban, agricultural, and marine foraging strategies, and demonstrated differences in dietary niche between age and colony cohorts.
5. Overall, DNA metabarcoding enabled the detection of a highly diverse diet and revealed differences in resource use among colonies and age cohorts. These findings demonstrate the potential of DNA metabarcoding to advance our understanding of diet in highly generalist species and contribute to ongoing efforts to understand how dietary variation may shape demographic processes in highly dynamic populations.

## Introduction

Spatial and temporal variation in animal diets provides key insights into resource use and availability and can help explain changes in individual fitness that scale up to population-level demographic shifts (Agashe & Bolnick, 2012; Griffin et al., 2022; O’Hanlon et al., 2017; Shutt et al., 2021). Quantifying individual diets of predators can also be used to estimate predation rates on native species or species of conservation concern (Robeson II et al., 2018; Shao et al., 2021; Shionosaki et al., 2015). However, despite the availability of several complementary methods for measuring diet (Hoenig et al., 2022; Lintulaakso et al., 2023), accurately quantifying individual diets remains challenging, as direct observation across an animal’s full foraging niche is rarely feasible.

The amplification and sequencing of DNA from faecal samples or from samples collected along the digestive tract, termed dietary DNA metabarcoding, is increasingly used to characterise animal diets and identify key prey (Alberdi et al., 2019; Williams et al., 2026). This approach can detect a broad range of consumed species that may be overlooked using conventional techniques, such as pellet and stomach contents analysis, and in addition does not necessarily require lethal sampling. However, DNA metabarcoding also poses significant methodological challenges, including amplification bias, contamination risk, and interference by host DNA, which can cause this approach to fail or yield biased results (Alberdi et al., 2019; Browett et al., 2021).

Despite their ecological importance across many ecosystems, generalist mesopredators present particular challenges for dietary metabarcoding (Tercel et al., 2021). This challenge, termed the ‘predator problem’ (Cuff et al., 2023), arises because metabarcoding relies on primer-based PCR amplification of target taxonomic groups that can amplify the DNA of both host and prey species. Most metabarcoding primers are designed to amplify broad groups such as eukaryotes, plants, or vertebrates, or narrower taxonomic groups (e.g., fish or mammals). When predators and their prey belong to different taxonomic groups, for example, piscivorous birds or mammals (Kennerley et al., 2024), insectivores (Mitchell et al., 2022; Verkuil et al., 2022), or herbivores (Hoff et al., 2025), this approach can be highly effective, as primers amplify dietary DNA rather than host DNA. In contrast, when both predator and prey fall within the same targeted taxonomic group, host DNA is often co-amplified at high proportions. This issue is particularly pronounced in systems where predators specialise on closely related taxa (e.g., sparrowhawks preying on other birds), but it also arises in studies of highly generalist predators where broad-range primers are required to detect diverse animal prey (Tercel et al., 2021). In such cases, host DNA can overwhelm dietary signal, making it difficult to recover informative diet data. This has contributed to a paucity of DNA-derived diet data for generalist mesopredators, a trophic group that is becoming increasingly common and plays a disproportionately important role in shaping ecosystem dynamics and the fate of vulnerable species, such as ground-nesting waders (Prugh et al., 2009).

The performance of DNA metabarcoding can depend on the type of biological sample used to characterise diet (Paprocki et al., 2024). Faecal samples are the most commonly used sample type (Ando et al., 2020), but others include cloacal, anal, pharyngeal or buccal swabs, regurgitated material, and, occasionally, swabs of the beaks and talons of raptors to detect prey DNA (Bourbour et al., 2019). Where carcasses are available, stomach contents and material from the lower digestive tract can also be analysed. These sample types vary in both the quantity of host DNA they contain and the degree of DNA degradation, factors that directly affect amplification success and taxonomic resolution. In faecal samples, prey DNA is typically highly fragmented as a result of digestive processes, which can reduce amplification efficiency and hinder reliable prey detection (Kamenova et al., 2018). Sampling methods, such as storage, can also influence the level of DNA degradation, which in turn affects amplification success (Graux et al., 2024).

One strategy to reduce the overwhelming amplification of host DNA is the use of blocking primers (Homma et al., 2022). These modified primers are designed to bind specifically to host DNA at the target locus and suppress amplification during PCR using a C3 spacer at the 3′ end to prevent extension. Blocking primers can substantially improve the recovery of dietary sequences when host DNA is abundant, although the extent to which they suppress amplification can vary. However, their use introduces additional challenges: they may bind non-target DNA, particularly from taxa closely related to the host, potentially biasing prey detection (Cuff et al., 2023). There is also some evidence that blocking primers can increase the relative amplification of trace contaminant DNA (Zabala et al., 2025), raising the risk of false positives. This concern may be especially relevant for low-biomass or highly degraded samples, such as faecal material, where prey DNA is already limited. Consequently, the trade-offs associated with blocking primer use likely vary across sample types, and evidence is needed on how best to balance their benefits and limitations in different settings.

In this study, we evaluated the performance of DNA metabarcoding using COI (targeting animal DNA) and 12S (targeting vertebrate DNA) primers for quantifying the diet of lesser black-backed gulls (*Larus fuscus*) at two breeding colonies in northwest England. Lesser black-backed gulls are highly generalist mesopredators and scavengers that consume a wide range of marine and terrestrial prey (Camphuysen et al., 2023; Coulson & Coulson, 2008; Gyimesi et al., 2016; Ingraham et al., 2020; O’Hanlon et al., 2025). Over the past 30 years, lesser black-backed gulls have undergone substantial changes in their ecology and distribution across the UK (Burnell et al., 2023), with widespread declines in monitored populations leading to their protected status (Burnell et al., 2023).

Some of the most dramatic shifts in lesser black-backed gull demography in the UK have occurred in northwest England. Traditional breeding strongholds, including South Walney Nature Reserve (Cumbria; hereafter ‘Walney’; Fig. 1a), have experienced severe declines. Breeding numbers at Walney fell from approximately 20,000 to just 380 pairs between 2000 and 2022. In contrast, breeding populations have increased at other sites, most notably the Bowland Fells region (Lancashire; hereafter ‘Bowland’; Fig. 1a), which now supports the largest lesser black-backed gull breeding colony in the UK (Burnell et al., 2023). This colony recovered from 4,000 pairs in 2012 to 31,000 pairs in 2023 (APEM, 2023), following the cessation of widespread culling that had been used to control the population for several decades until the early 2000s (Burnell et al., 2023; Mitchell et al., 2004; Ross-Smith et al., 2014). Population estimates from the 1970s prior to culling range around 20,000 pairs (Duncan, 1981; Lloyd et al., 1991; Wanless & Langslow, 1983). Although these demographic shifts will be driven by multiple factors, resource availability and use around breeding colonies, particularly during chick rearing, are expected to play a central role.

Demographic shifts in mesopredator populations, including gulls, can also have cascading effects on other species through changes in predation pressure. Recent growth of the Bowland colony has increased concerns about potential impacts on breeding waders, whose conservation status has declined in recent decades (Sim et al., 2005). Large gull species are opportunistic predators (Ingraham et al., 2020) and some individuals may specialise on particular prey, occasionally exerting substantial effects on prey populations (Sanz-Aguilar et al., 2009). The areas in and around Bowland Fells provide important breeding habitat for lapwing (*Vanellus vanellus*), curlew (*Numenius arquata*), and oystercatcher (*Haematopus ostralegus;* McGuire, 2024), although the extent to which gulls predate these vulnerable species remains uncertain. Diet metabarcoding may therefore provide valuable insights into both the breadth and variability of lesser black-backed gull diets across areas experiencing contrasting population trajectories, while also helping to assess the potential consequences of demographic change for other species. Ultimately, this can provide evidence to better inform management and decision-making in these landscapes.

Within this context, our specific objectives were to:

1. Compare the performance of two primer sets (COI and 12S) for diet analysis across different sample types collected from lesser black-backed gulls, and evaluate whether three gull-specific blocking primers improve diet detection in 12S-amplified samples.
2. Assess the use of pharyngeal swabs to detect key species of conservation concern.
3. Describe overall gull diet composition and diversity.
4. Examine differences in dietary composition between age groups and colony cohorts.
5. Identify key indicator species that may suggest different foraging strategies.

To do this, we collected different sample types from pre-fledging lesser black-backed gull chicks during ringing at both breeding sites. We also sampled adult and sub-adult lesser black-backed gull carcasses of birds taken under licence for the conservation of ground-nesting waders in Bowland and submitted for scientific research. These carcasses were dissected to examine stomach contents for hard parts, with a particular focus on identifying wader remains. Swab samples were collected from the stomach, intestines, cloaca, and pharyngeal tract for DNA metabarcoding.

## Methods

### Sample collection from chicks for DNA metabarcoding

Faecal and regurgitate samples were collected during yearly colour ringing of lesser black-backed gull chicks (∼28 days old) across Bowland in early July 2023 and 2024 (n = 52 regurgitate samples; n = 58 faecal samples from 101 chicks), with paired samples from the same individual generally not possible. We also analysed a smaller number of faecal and regurgitate samples from chicks at South Walney, Cumbria, collected in late June and early July 2024 (n = 9 faecal samples; n = 7 regurgitate samples from 15 individuals). Faecal and regurgitate matter were sampled with sterile swabs from clean plastic holding containers that were disinfected after each use, or during ringing, taking care not to touch the container or other surfaces. Regurgitation occurs as an anti-predator response during handling, and regurgitates that are not well digested can often be assessed visually for content, yet are often too digested for visual analysis. Swabs were put into 2.0 mL microtubes filled with 1.0 mL of RNA/DNA Shield (Zymo) to prevent DNA degradation. Samples were stored at -20 °C for long-term storage until DNA extraction.

### Sample collection and stomach contents analysis of lesser black-backed gull carcasses

We dissected 107 adult lesser black-backed gull that were legally taken under licence for the protection of wader chicks and nests and submitted for scientific research. Birds were taken from May to July 2024 in the region around Bowland in areas with high breeding wader densities, and frozen within 24 hours postmortem. Based on gonad analysis and biometrics, the majority (>60%) of birds were male, with adult (>5 year old) plumage. Digestive tract sampling and stomach-contents (‘hard parts’) examination followed published sources (Carss, 2022; Marquiss et al., 1998). The full dissection and stomach-contents analysis method for fish-eating birds is described by Carss et al. (2012) for fish-eating birds was adapted slightly for this lesser black-backed gull study. After thawing, the body cavity was opened, following the digestive tract from the beak to the cloaca. Any whole, intact, undigested prey – as well as any partially-digested material – present in the proventriculus/gizzard was examined and measured (to the nearest millimetre) and then subsequently stored refrozen. Examination of prey remains was confined to hard parts in the stomach contents material. Bird-derived hard parts in the stomach contents were cleaned, examined, and identified where possible. Feather colouration and growth stage were noted, and biometric measurements taken whenever feasible. Species identification of prey were based on these characteristics and published guides (Harrison, 1980).

For all 107 birds, proventriculus/gizzard swabs and intestinal swabs (from four incisions along the length of the intestine) were collected. Where stomach contents yielded multiple potential prey items, specific prey items from the proventriculus/gizzard were swabbed individually. All swabs were placed in 1ml of RNA/DNA Shield (Zymo) buffer for DNA preservation and frozen at -20°C. All samples were collected using sterile latex gloves, gloves were changed between each dissection, all surfaces were cleaned with bleach (containing 4.5g sodium hypochlorite per 100g) diluted by a factor of 10 diluted bleach, and dissection equipment was sterilized with diluted bleach between samples. Scalpels and other dissection tools were soaked in bleach and rinsed between both dissections and sampling steps. At the start of each day, the lab bench, sample tray, dissection equipment, and gloved hands were swabbed to act as a negative sampling control (see section on negative controls).

It is important to note that lesser black-backed gull adults and chicks were sampled at different times of year, and that adult sampling specifically targeted individuals foraging within breeding wader habitat. The adult dataset is therefore biased towards individuals using this foraging strategy, and may not be representative of the wider adult population.

**Figure 1).**
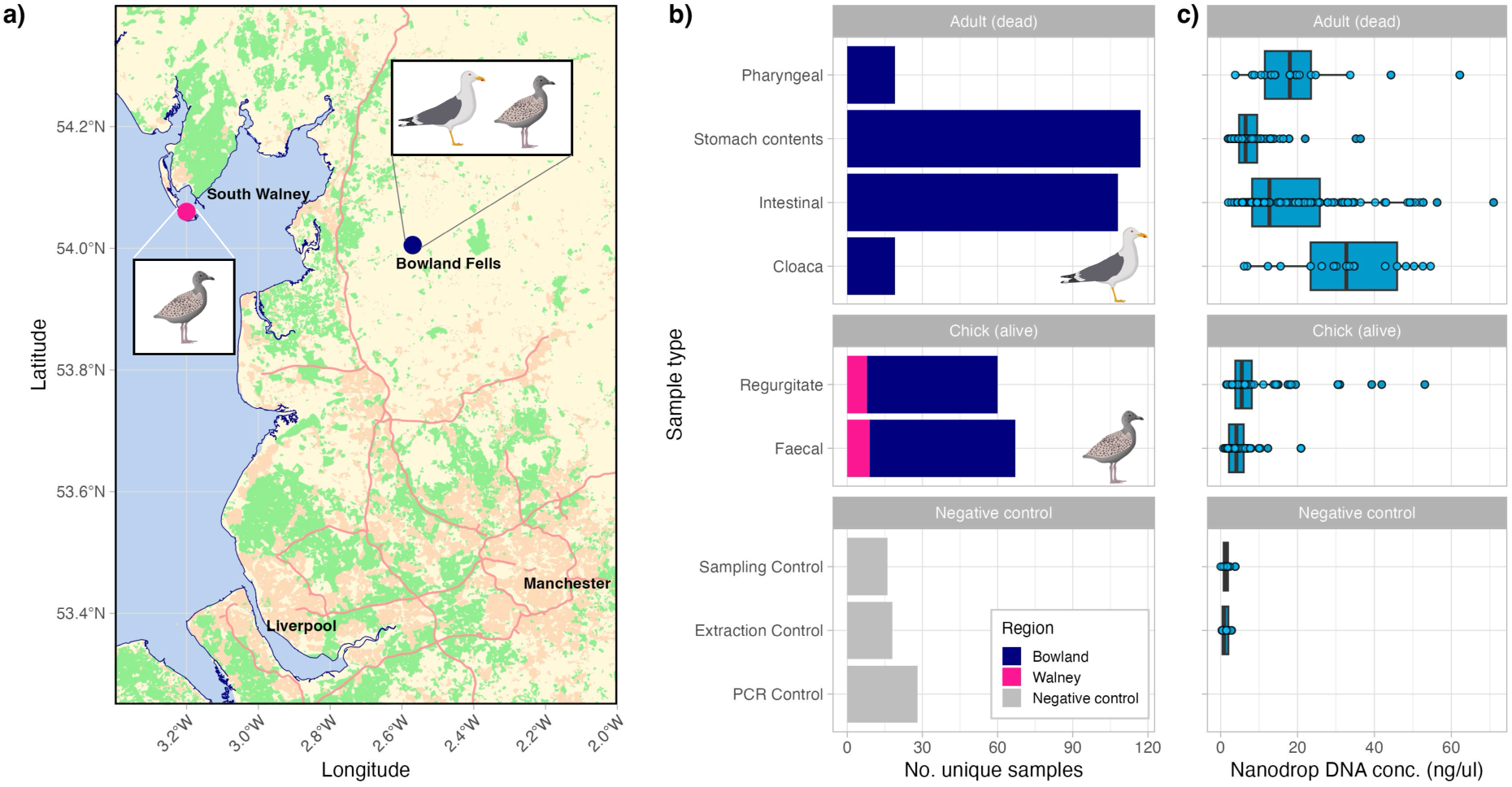
Sampling locations, sample sizes and DNA concentrations of samples used in this study. (a) Map highlighting Walney and Bowland where gulls and samples originated; b) Number of different sample types collected from Bowland (blue) and South Walney (pink), and negative controls (grey; note the 28 PCR controls are not included as these are not ‘samples’ that could be quantified with the Nanodrop); c) Approximate DNA concentration of samples using Nanodrop. Map data downloaded from Open Street Maps.

### DNA extraction

All samples were extracted using the ZymoBIOMICS DNA Miniprep Kit (Zymo), following the manufacturer’s protocol with minor modifications. Samples, stored in DNA/RNA shield buffer (Zymo), were initially vortexed, and DNA extractions were conducted using 300 μL of the buffer with 400 μL of buffer AL added to the mixture. All other steps followed the kit protocol. DNA extractions were quantified using a Nanodrop spectrophotometer before DNA amplification. One negative extraction control was included for every extraction batch, consisting of a maximum of 12 samples and totalling 18 DNA extraction controls (see section on negative controls).

All samples were processed in a dedicated pre-PCR laboratory free from amplified PCR products. Personnel wore appropriate personal protective equipment at all times, and all surfaces and equipment were routinely decontaminated with 50% bleach followed by 70% ethanol. Gloves were changed frequently and disposed of after use. Filtered pipette tips were used throughout all procedures to minimise the risk of contamination. DNA concentration was estimated using Nanodrop, with unusually high concentrations of DNA found in intestinal and cloacal samples (Fig. 1c).

### Amplification primers and blocking primer design

In this study, we used two different primer sets targeting ∼106 bp of the rRNA 12S mitochondrial gene (Kelly et al., 2014; Riaz et al., 2011) and ∼313 bp of the mitochondrial COI gene (Leray et al., 2013, Geller et al., 2013) genes for amplification of all vertebrate and metazoan (animal) rRNA genes, respectively (Table 1a). These primers were chosen in order to capture the broad range of vertebrate and invertebrate food items consumed by gulls.

Initial testing of COI and 12S primer sets on gull faecal samples shows that both primers largely yielded gull DNA in these samples. We therefore designed and tested three potential blocking primers to inhibit the amplification of gull DNA during 12S amplification (Table 1b). We did not design blocking primers for the COI primer set owing to its high degree of degeneracy. To design blocking primers targeting gulls, multiple sequence alignments were prepared using 55 sequences of the 12S ribosomal RNA region, comprising 38 bird species representing genera known to occur in the UK (McInerny et al., 2022), including lesser black-backed gull (Table S1), and 15 species of non-bird vertebrates expected to be detected (e.g., *Bos taurus*, *Gadus* spp.) using the 12S-V5 primer set (Riaz et al., 2011).

Both target (lesser black-backed and other gulls) and non-target sequences were aligned using Clustal W in MEGA Software (Kumar et al., 2024). The alignment process aimed to identify regions where the blocking primer would bind, typically targeting areas where sequences overlap. Primers were then designed using Geneious Prime, with the melting temperature (Tm) calculated using OligoCalc (http://biotools.nubic.northwestern.edu/OligoCalc.html) based on the sequence composition. For all designed blocking primers, to prevent elongation by the polymerase, a 3′-spacer C3 modification was incorporated into the blocking primer.

The functionality of the blocking primer was manually validated in silico using Basic Local Alignment Search Tool (BLAST), ensuring its effectiveness in blocking the amplification of gull DNA sequences. When checking the designed blocking primers, BLAST returned only matches corresponding to Laridae species (e.g., *Larus* spp., *Pagophila eburnea*, *Rissa tridactyla*), demonstrating the blocking primer efficiency in binding to gull DNA sequences.

Three different types of blocking primers were designed, including one annealing inhibiting blocking oligo (i.e. prevents amplification of metabarcoding primers due to the presence of blocking oligos in the binding sites; BLK1), one elongation arrest blocking oligo (i.e., binds to a region near the metabarcoding primer site, physically preventing the polymerase from synthesizing the DNA strand further; BLK2) and one Dual Priming Oligo (DPO; BLK3).

DPO consists of two separate priming regions linked by a polydeoxyinosine linker, designed to reduce the melting temperature (Tm) of long primers and prevent mispriming, primer-dimer, and hairpin formation. DPOs are particularly useful when it is difficult to find a suitable binding site for a blocking primer adjacent to a metabarcoding primer’s binding site. The primer includes two segments connected by five deoxyinosines and a C3 spacer modification at the 3′ end.

Blocking primer tests were conducted, including stomach content, intestinal, faecal and regurgitate samples. Following blocking primer optimisation experiments, which included testing different blocking primer concentrations (results presented below), BLK2 at a 10x concentration was selected and incorporated into all 12S amplification reactions across sample types.

**Table 1).**
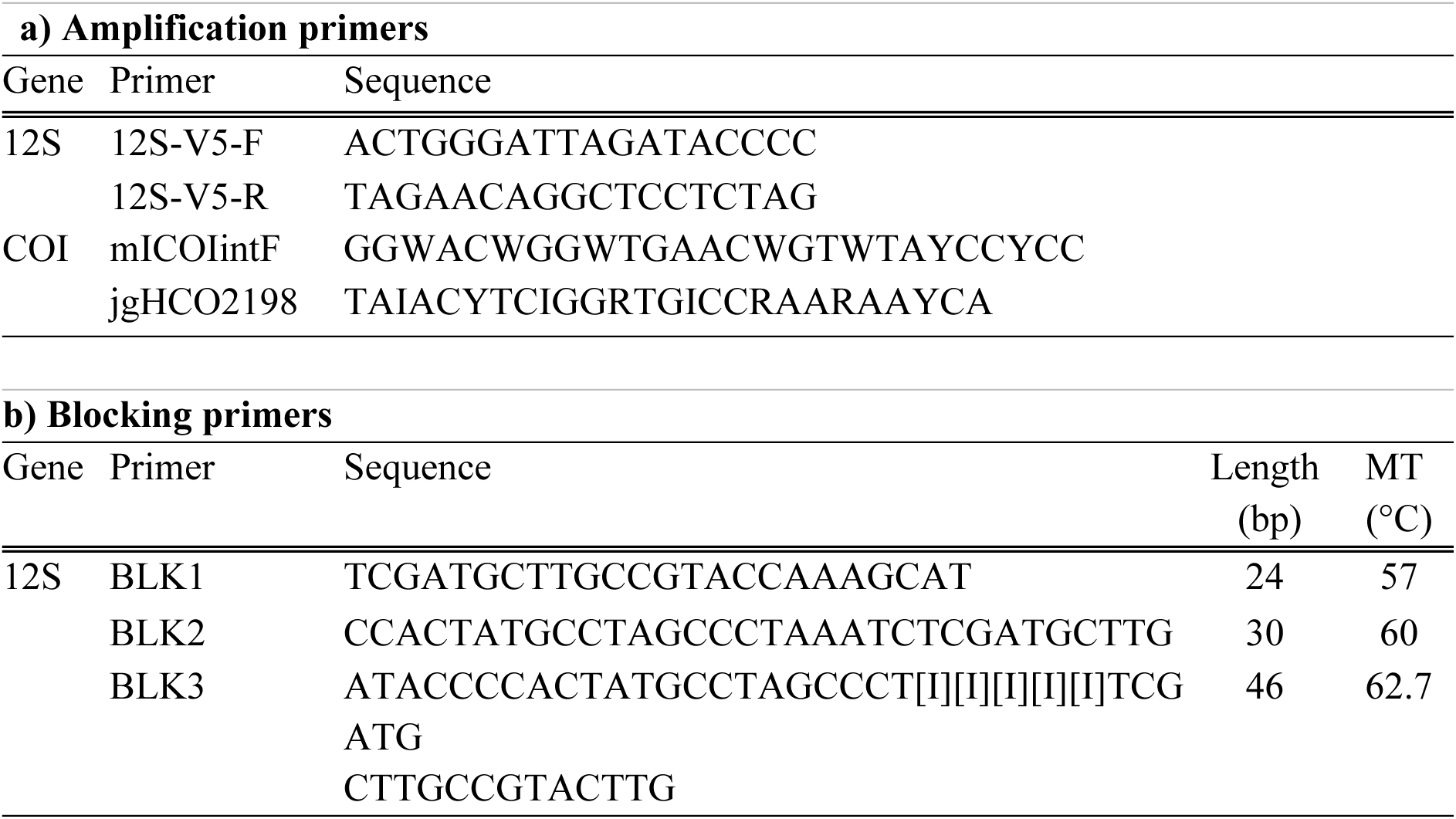
Amplification and blocking primers tested, including sequences, primer length in bp and melting temperatures (°C).

### DNA amplification

PCR reactions were prepared in triplicate to account for amplification stochasticity, and each primer pair included unique eight-base multiplex identifier (MID) tags to enable sample multiplexing prior to library preparation. PCR reactions were carried out in 20 μL reactions, containing 10 μL AmpliTaq Gold™ 360 Master Mix (Applied Biosystems), 0.16 μL bovine serum albumin (ThermoScientific), 1 μL of each primer (5 μM), ultra-pure water (adjustable volume), blocking primers, and 2 μL of DNA template. For 12S, reactions containing blocking primers were adjusted according to the desired metabarcoding: blocking primers ratio.

For 10x blocking primer concentration, PCR reactions were carried out in 20 µL volumes, containing 10 µL of AmpliTaq Gold™ 360 Master Mix (Applied Biosystems), 0.16 µL of bovine serum albumin (ThermoScientific), 2 µL of Riaz primers (5 µM), 1 µL of blocking primer BLK2 (100 µM), 4.84 µL of ultra-pure water, and 2 µL of DNA template. Thermal cycling conditions for COI were: initial denaturation at 95°C for 15 min; 10 cycles of 94°C for 30 s, 49°C for 45 s, and 72°C for 30 s; followed by 30 cycles of 95°C for 30 s, 47°C for 45 s, and 72°C for 30 s; and a final extension at 72°C for 10 min. For 12S rRNA, thermal cycling conditions were: initial denaturing step of 95°C for 5 min, 35 cycles of 95°C for 15s, 57°C for 30s, and 72°C for 30s and a final extension step of 60°C for 30 min. All PCR products were verified by electrophoresis on a 1.5% agarose gel.

Negative PCR controls were included on all PCR plates (see section below on negative controls). PCR negative controls were generated by adding nuclease-free water in place of template DNA to monitor potential contamination introduced during amplification. In addition, 5–7 sampling and extraction controls relevant to each sample batch were incorporated into the same plate.

### Negative controls

To detect, quantify, and account for contamination, we generated metabarcoding data for 62 negative controls comprising three control types: sampling controls (n = 16; one per day of dissections), extraction controls (n = 18; one per extraction batch), and PCR controls (n = 28; 4–6 per sequencing plate). Sampling controls were used to detect contamination introduced during bird handling and sample collection, extraction controls identified contamination arising during DNA extraction (e.g., reagent contamination or cross-contamination between samples), and PCR controls assessed contamination introduced during library preparation and amplification. All negative controls were processed identically to biological samples, including the addition of the 12S blocking primer. The blocking primer suppresses amplification of gull DNA and consequently increases amplification sensitivity for low-level contaminant DNA (Figs. 3 and 4). Together, these controls allowed contamination to be assessed at multiple stages of sample processing.

Two positive controls consisting of *Brachyteles hypoxanthus* DNA (the northern muriqui, a non-native primate species) were included on the first two plates to assess potential cross-contamination during PCR, but, due to positive controls increasing overall contamination risk, were deemed unnecessary for further plates.

### DNA sequencing

Upon PCR amplification, samples were pooled into four libraries (one per plate) per primer set, totalling eight libraries. A left-sided size selection was performed using 1.2× Agencourt AMPure XP beads (Beckman Coulter; total of 800 μL used), and library preparation was carried out using the KAPA HyperPrep Kit (Roche) with dual-indexed adapters. Libraries were normalised and sequenced at Novogene using the Illumina NovaSeq X platform.

Sequencing was conducted using partial lane allocation alongside other projects, targeting a minimum of 6 Gb of raw data per library, with a 1% PhiX spike-in applied to low-diversity libraries. The 12S libraries were sequenced using 2 × 150 bp paired-end reads, and the COI libraries using 2 × 300 bp paired-end reads.

### Bioinformatics

Bioinformatic analyses were performed using the OBITools 1.2.2 metabarcoding package (Boyer et al., 2016). Read quality was first assessed with FastQC (v 0.12; Andrews, 2010). Paired-end reads were then aligned using the Illumina paired end command (score>40.0), followed by demultiplexing and primer removal using *ngsfilter*. Size selection was conducted with *obigrep* (seq_length 80-150 bp), which removed fragments likely resulting from library preparation artefacts (e.g., primer-dimers, non-specific amplifications) as well as reads containing ambiguous bases.

Unique sequences were subsequently clustered and chimaeras removed using VSEARCH v2.0 (Rognes et al., 2016), applying the uchime-denovo algorithm (Edgar et al., 2011).

Sequences were then grouped into Molecular Operational Taxonomic Units (MOTUs) using SWARM v3 (d = 1; Mahé et al., 2021). Singletons were excluded, and the remaining MOTUs were queried against the GenBank nucleotide database using BLASTn. For taxonomic assignment, sequences required a minimum of 90% alignment and over 80% similarity to be considered a valid match. The top 25 BLAST hits were retrieved, and the most frequently occurring taxonomic identifier (taxid) was assigned. MOTUs showing ≥98% identity and matching a single species were assigned at the species level (Arrizabalaga-Escudero et al., 2018; Clare et al., 2014). MOTUs with identity between 95–98% or matching multiple species were assigned at the genus level. Those with 93–95% identity were assigned to family, and those between 90–93% to order. To enhance taxonomic resolution, MOTUs with low-confidence assignments were manually cross-checked using the BOLD database.

### Data processing

#### MOTU count correction

MOTUs were agglomerated to species-level taxonomy where species-level assignments were available; these are referred to as taxa hereafter. To account for low-level contamination, read counts were corrected separately for each sequencing plate using the associated negative controls. For each taxon, the maximum read count observed across all negative controls on a plate was subtracted from the corresponding read counts in all samples processed on that plate, as is recommended best practice (Drake et al., 2022).

#### Filtering

Following count correction, taxa with ≤100 corrected reads were removed from the dataset. Taxa not assigned to Animalia or Metazoa were also excluded. Additional filtering was then applied to remove known contaminants identified from the negative controls (e.g, human DNA and non-target taxa associated with previous laboratory/sequencing work).

Primer-specific contaminants detected only in COI libraries were removed from the COI dataset only. The COI negative controls contained contaminating MOTUs assigned to several commercial fish species, including *Pollachius virens* (pollock), *Scomber scombrus* (Atlantic mackerel), *Lophius piscatorius* (anglerfish), *Pangasius bocourti* (basa catfish), *Gadus morhua* (Atlantic cod), and *Conger conger* (European conger eel). These taxa were primarily detected in sampling controls rather than extraction or PCR controls, suggesting contamination during sample collection or storage rather than amplification. Because these MOTUs were largely absent from 12S controls despite being amplifiable with 12S primers, they were considered primer-specific contaminants and removed from the COI dataset.

Because these species are also amplifiable with the 12S primers, any genuine occurrence in the samples would be expected to be detected in the 12S dataset, minimizing the risk that their removal from the COI dataset would exclude true biological detections.

#### Assignment of diet categories

Detected taxa were assigned to broad dietary categories corresponding to their likely origin, including urban-sourced animal products, commercial and non-commercial fish, and wild mammals and birds. The presence of non-commercial fish taxa, such as flounder and pipefish, was interpreted as evidence of marine foraging, whereas the presence of certain livestock species, such as chicken, was used as an example considered as an evidence of urban-sourced animal products.

#### Data analysis and statistics

Phylogenetic relationships between prey species was generated using the Open Tree of Life and rotl R package (Michonneau et al., 2016). To test for significant differences in prey diversity detected between sample types, we fitted a General Linear Model with a gaussian distribution using the function *glmmTMB::<u>glmmTMB()</u>*. For analysis of dietary niche between the three cohorts (Bowland adults, Bowland chicks, Walney chicks) we agglomerated detections by individual. To identify clustered of correlated diet items, we performed a Spearmans correlation on pairwise detections, including only species that occurred at least three times. We then created a cluster-based heatmap applying *heatmap::pheatmap()* using Euclidean distances, and built a network using the igraph package. To identify differences in dietary niche between cohorts, we applied a constrained ordination on Jaccard Distance, using the *vegan::capscale()* function in R, and tested for significance using a PERMANOVA. All data processing and analysis was conducted in R (version 4.4).

## Results

### Proportion of host and human DNA across sample types without blocking primers

Overall, regurgitate samples had the lowest proportion of host reads for both 12S and COI amplification primers, whilst faecal and intestinal samples had the highest proportion (Fig. 2a). For 12S primers, faecal and intestinal samples were almost entirely composed of gull reads (faecal samples: 96.5 ± 7.6 %; Intestinal samples: 99.7 ± 0.4 %), whereas regurgitate and stomach-content samples were more variable but averaged around 60–65% gull DNA (regurgitate samples: 60.4 ± 42 %; Stomach content samples: 65% ± 34 %). For COI primers, proportion of gull reads was variable across all sample types, with higher proportions of gull reads in faecal (89 ± 19 %), intestinal (97 ± 9 %), and stomach content samples (85 ± 23%), and lower proportions in regurgitate samples (55 ± 33 %).

The proportion of contaminating human reads was very low for all sample types when processed without blocking primer (<0.1%; Fig. 2b).

### The performance of blocking primers at reducing host DNA across sample types

We tested three different blocking primers to reduce host DNA in 12S-amplified samples. An ideal blocking primer should not only reduce host DNA but also avoid biasing results by blocking non-target taxa or amplifying contaminants.

In terms of blocking host DNA, each of the three blocking primers performed differently depending on sample type (Fig. 2a). Blocking primer 1 (BLK1) did not effectively reduce host DNA across any sample types and was therefore not considered further.

Blocking primer 3 (BLK3) was the most effective at blocking gull DNA across all sample types (Fig. 2a). However, it also increased the amplification of contaminating reads, with human DNA contamination being an average of ∼1% in faecal and intestinal samples, but sometimes reaching 30% (Fig. 2a). It also appeared to inhibit the detection of prey taxa confirmed through stomach content analysis (e.g., lapwing). BLK3 performance was also highly dependent on concentration, with – counterintuitively – lower (5x) concentrations being more effective at reducing gull DNA (Fig. S1a) but also inflating contaminating DNA more (Fig. S1b).

Blocking primer 2 (BLK2) only slightly reduced host DNA in faecal samples compared to controls (Fig. 2a) but had no noticeable effect on host DNA in intestinal samples or regurgitate samples. It also increased the amplification of contaminant DNA, although to a lesser extent than BLK3 (Fig. 2b). Although the reduction in gull DNA in faecal samples was modest (from 97% down to 89%), it was sufficient to allow improved detection of dietary DNA. Visual inspection of the broader dataset suggested that BLK2 did not inhibit detection of target prey species, such as lapwing, unlike BLK3.

Based on these results, BLK2 was selected as the most effective blocking primer for inclusion in subsequent 12S PCR reactions. All 12S results presented hereafter were generated using the BLK2 blocking primer.

**Figure 2).**
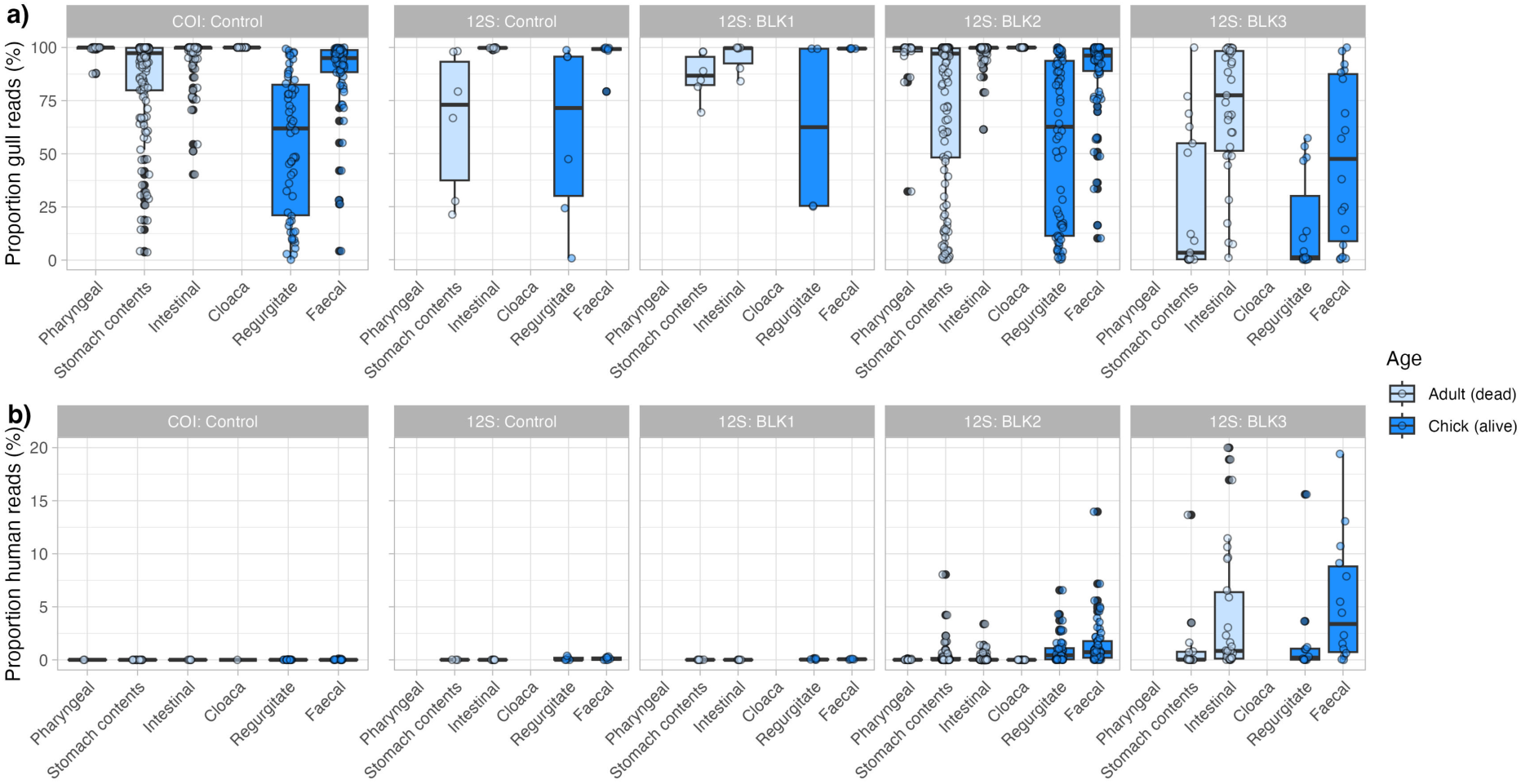
Performance across sample types, primers, and blocking primers tested; a) The proportion of gull reads across tested sample types; b) the proportion of contaminating human reads across tested sample types. Not all blocking primers were tested on all sample types, and these are indicated with ‘no data’. For clarity, we indicate which sample types were sourced from chicks and dead adults.

### Diet composition and diversity across sample types and amplification primers

In total, we detected 71 unique species belonging to 61 genera, including 38 species of vertebrates, 15 species of arthropods, 13 species of annelids, and 5 species of molluscs (Fig. 3). The most frequent species identified were *Bos taurus* (cattle, including from consuming beef and dairy), *Ovis aries* (sheep, mutton), *Sus scrofa* (pig, pork), *Gallus gallus* (chicken), *Vanellus vanellus* (European lapwing), and earthworms of the genus *Lumbricus* and *Aporrectodea*, especially *L. terrestris*. *Pleuronectes platessa* (European plaice) was also relatively common in birds sampled at Walney.

**Figure 3).**
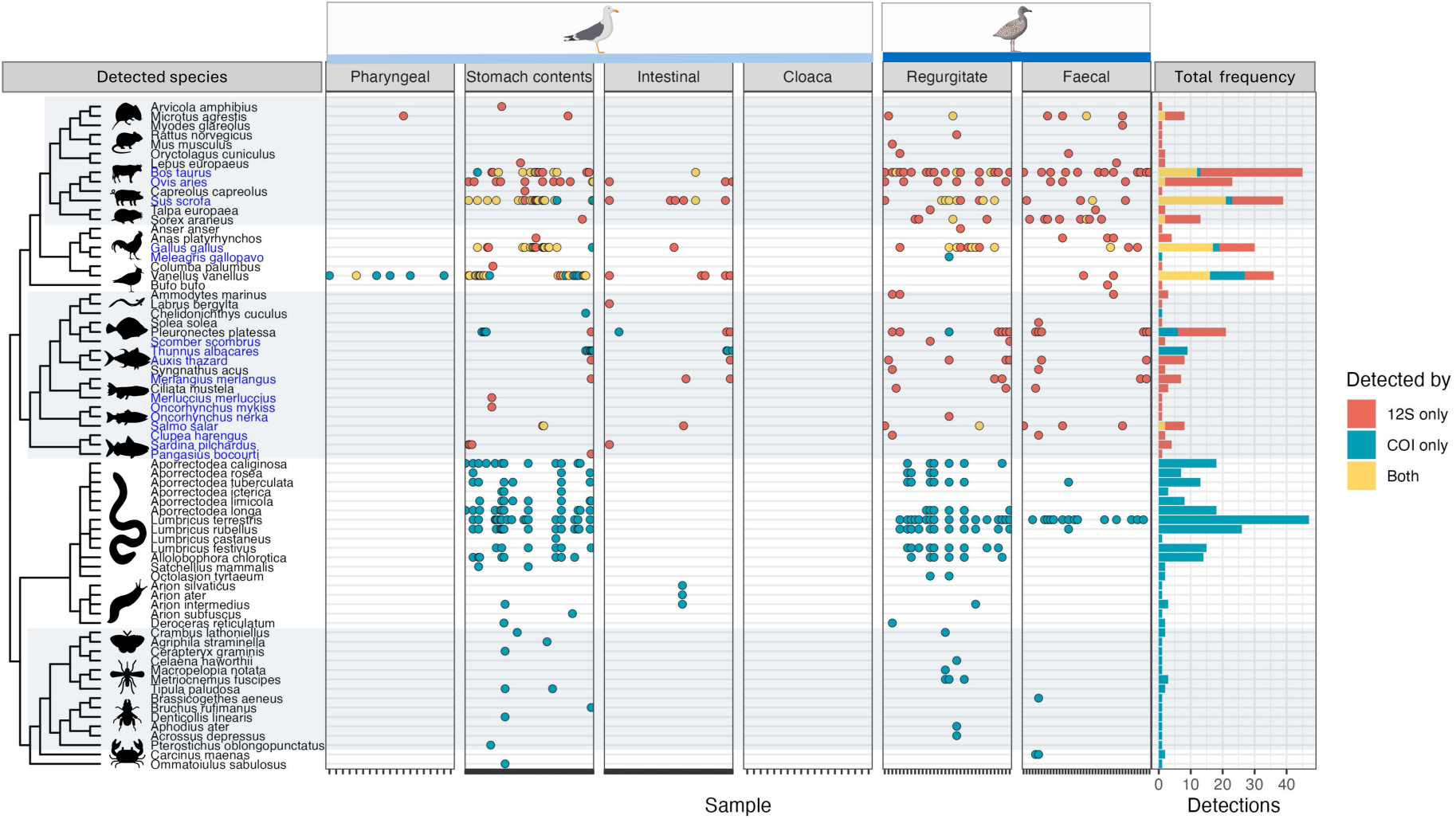
Prey detections across samples, facetted by sample type. Prey taxa are arranged on the y-axis according to their phylogenetic relationships, and commercial species are coloured blue. Detections (dots) are coloured according to whether they were detected by 12S primers only, COI primers only, or both 12S and COI. The panel on the right shows the frequency of detections for each primer set. Light blue panels highlight mammal species, fish, and arthropod species.

Regurgitate samples yielded the highest diversity of prey species (mean = 5, max = 15), whilst faecal samples (with blocking primer) detected an average of two and maximum of six species per sample (Fig. 4; see Table S2 for GLM). Despite not detecting as many species as regurgitate samples, faecal samples overall detected a similar suite of species at the cohort level compared to regurgitate samples (Fig. 3). In contrast, intestinal and cloacal swabs collected from dead adults performed poorly across both primer pairs, even when blocking primers were used (Fig. 3, Fig. 4), suggesting that DNA in these sample types was highly degraded or fragmented.

Comparisons between chick faecal and regurgitate samples also revealed clear differences in primer performance between these sample types. 12S primers detected a similar diversity of prey species across sample types (regurgitate = 22 species; faecal = 21 species). However, COI primers performed poorly on faecal samples (regurgitate = 25 species; faecal = 9 species). In these samples, the COI primers consistently detected only a single earthworm species, *Lumbricus terrestris*, despite at least eight earthworm species being commonly identified in regurgitate samples and numerous vertebrate taxa being detected by the 12S primers (Fig. 3). This difference may reflect the longer fragment amplified by the COI primers, which could reduce amplification success when prey DNA is highly fragmented, as is typical of faecal material.

Overall, both 12S and COI primers are designed to detect vertebrate taxa, vertebrate species were much better represented in samples amplified with the 12S primers (including the blocking primer) than with the COI primers. In contrast, the COI primers primarily detected annelids, particularly earthworms and slugs (Fig. 3). Despite this, both primer sets detected lapwing DNA in stomach content swabs (Fig. 3), likely because these samples contained a relatively large amount of prey material.

**Figure 4).**
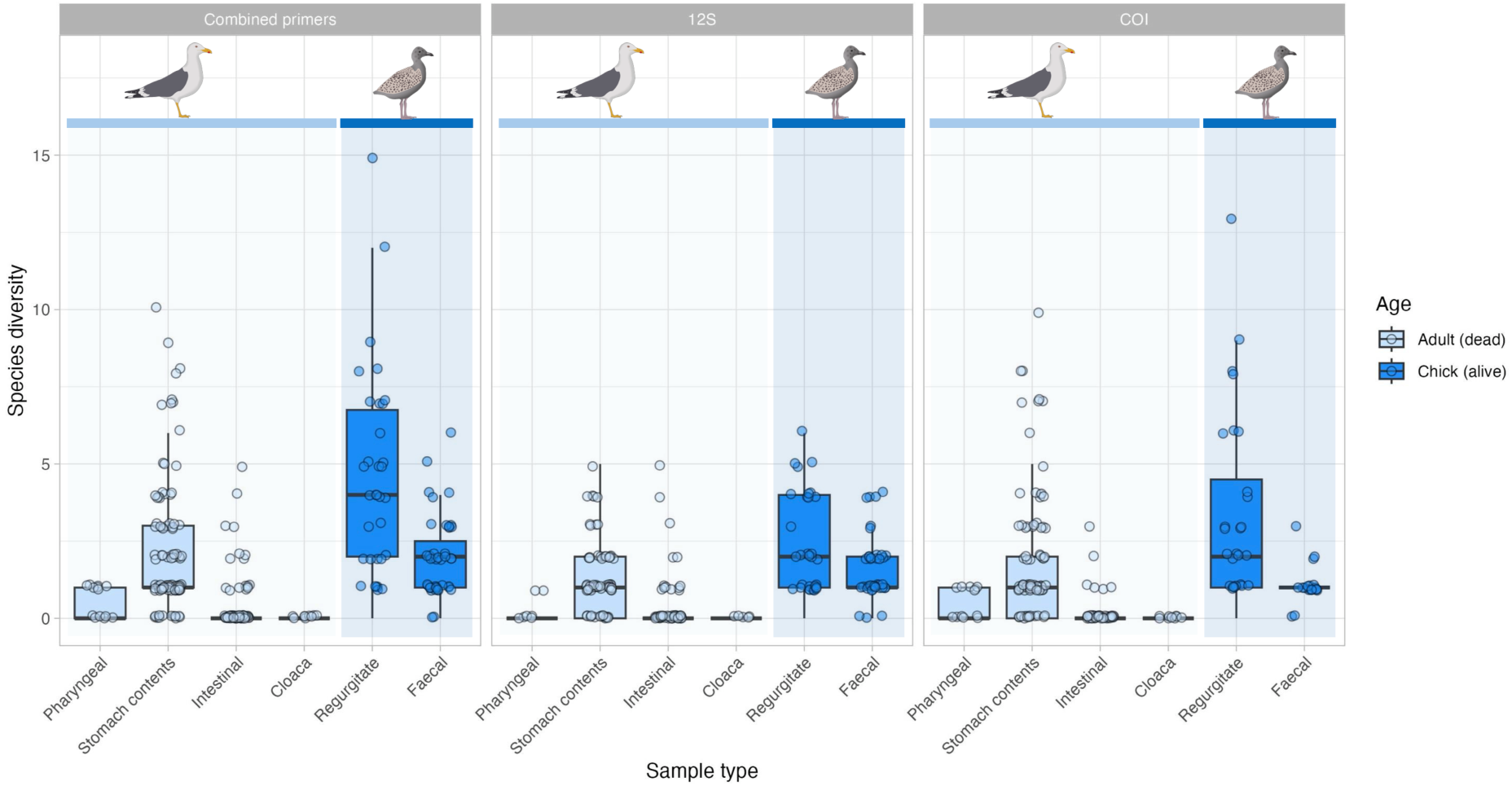
A comparison of species diversity detected by 12S and COI primers (and combined) across different sample types collected from lesser black-backed gulls. For clarity, we indicate which sample types were sourced from either chicks or adult carcasses.

### Pharyngeal and stomach swabs detect recently consumed key prey

Overall, DNA metabarcoding of stomach-content swabs, which contained recently consumed prey remains, aligned well with visual inspection and provided additional taxonomic resolution for prey items that could not be identified morphologically (Table S2). In total, 15% of dissected adult gulls contained either European lapwing (*V. vanellus*) remains or unidentified wader remains in their stomachs. DNA metabarcoding identified all of these wader remains as European lapwing and additionally detected lapwing DNA in the stomachs of three further birds. Almost all lapwing detections occurred in birds collected during May, with only a single lapwing detection in a bird collected in July. Small-mammal teeth recovered from the stomach of one bird were identified as the short-tailed field vole (*Microtus agrestis*). Twenty-one birds had empty stomachs at the time of dissection. Of these, trace DNA from earthworms and urban-associated food items, such as chicken, was detected in nine individuals, whereas no DNA was detected in the remaining birds (n = 12). Overall, DNA metabarcoding provided additional dietary information but likely captured only recently consumed prey, rather than the full diet accumulated over several days.

We assessed whether pharyngeal swabs could be used to detect key prey species that were also present in the stomach. Pharyngeal samples were collected from 19 adult birds that were subsequently dissected. Eight of these individuals contained confirmed lapwing remains in their stomachs, as determined by both dissection and DNA metabarcoding of stomach-content swabs. Lapwing DNA was detected in pharyngeal samples from six of these eight birds (75%), with no false-positive detections. The only other vertebrate prey item that could be confidently identified from stomach contents was a Short-tailed field vole (*M. agrestis*; *n* = 1), which was also detected in the corresponding pharyngeal sample. This finding suggests that pharyngeal swabbing may be capable of detecting prey several hours after consumption. Notably, lapwing detections from pharyngeal swabs were obtained almost exclusively using COI primers, with few or no detections using 12S primers.

### Dietary niches of different age and colony cohorts

To investigate potential differences in diet composition and dietary niche among lesser black-backed gull age classes and breeding colonies, we compared diets across three sampled cohorts: Bowland adults (sampled May–July), Bowland chicks (sampled July), and Walney chicks (sampled July; Fig. 5). For these analyses, DNA metabarcoding detections from different primer sets and sample types were aggregated at the individual level, providing a single dietary profile for each bird.

Across sampled gulls, the most frequent diet items were cow (*B. taurus*), earthworm species (especially *L. terrestris* and *L. rubellus*), European plaice (*P. platessa)*, and pig (*S. scrofa*) and chicken (*G. gallus;* Fig. 5). However, the proportions of different diet items differed between cohorts. Bowland adults and chicks showed frequent detections of other urban-associated food items, including pig (*S. scrofa*) and chicken (*G. gallus*). They also exhibited a greater diversity of earthworm taxa, alongside detections of European lapwing in adults and small mammals in chicks. In contrast, although sample sizes were smaller at Walney, chick diets appeared to contain a higher proportion of marine prey, with fish detected frequently and European plaice being particularly common.

**Figure 5).**
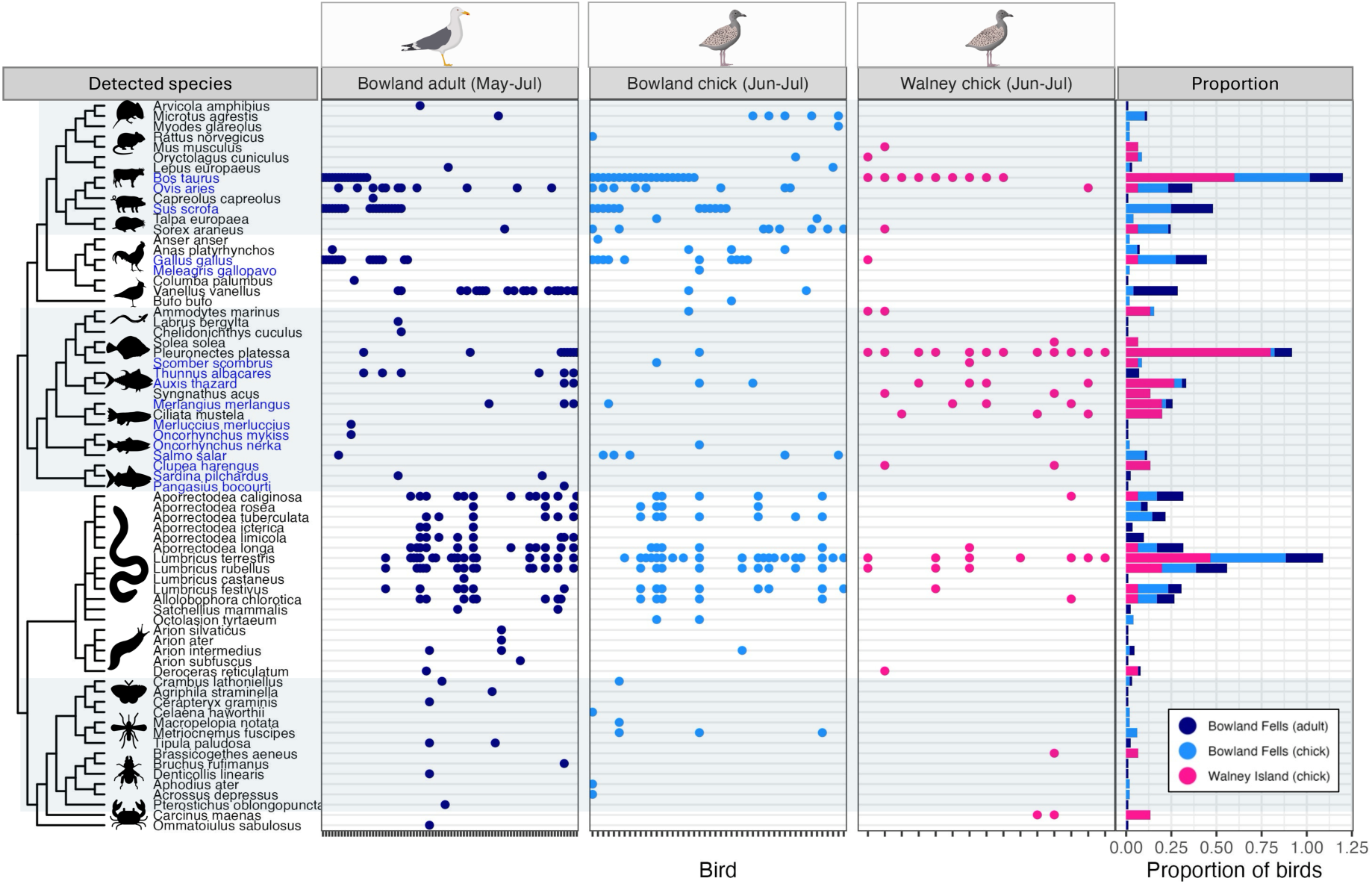
Prey detections across individual birds, facetted by age and colony cohort. Prey taxa are arranged on the y-axis according to their phylogenetic relationships, and commercial species are coloured blue. The panel on the right shows the proportion of detections per cohort. As sample sizes differ between cohorts, we present proportions rather than frequency. Light blue panels highlight mammal species, fish, and arthropod species.

A constrained ordination, a method designed to detect key differences between cohorts, revealed significant differences in dietary niche between cohorts (PERMANOVA; F = 4.0, p < 0.001), driven by specific dietary taxa (Fig. 6a). Walney chicks were characterised primarily by European plaice and, to a lesser extent, other fish species, including the coastal fish fivebeard rockling (*Ciliata mustela*). In contrast, Bowland chicks were characterised by the earthworm *L. terrestris*, pig, and the common shrew. A small number of Bowland adults clustered with Walney chicks, reflecting a more coastal or marine diet, yet this cohort was distinguished from other cohorts by a greater prevalence of lapwing in the diet (likely reflecting sampling bias from targeting gulls in breeding wader habitat) and the earthworm *Aporrectodea limicola*, a wet soil specialist.

**Figure 6).**
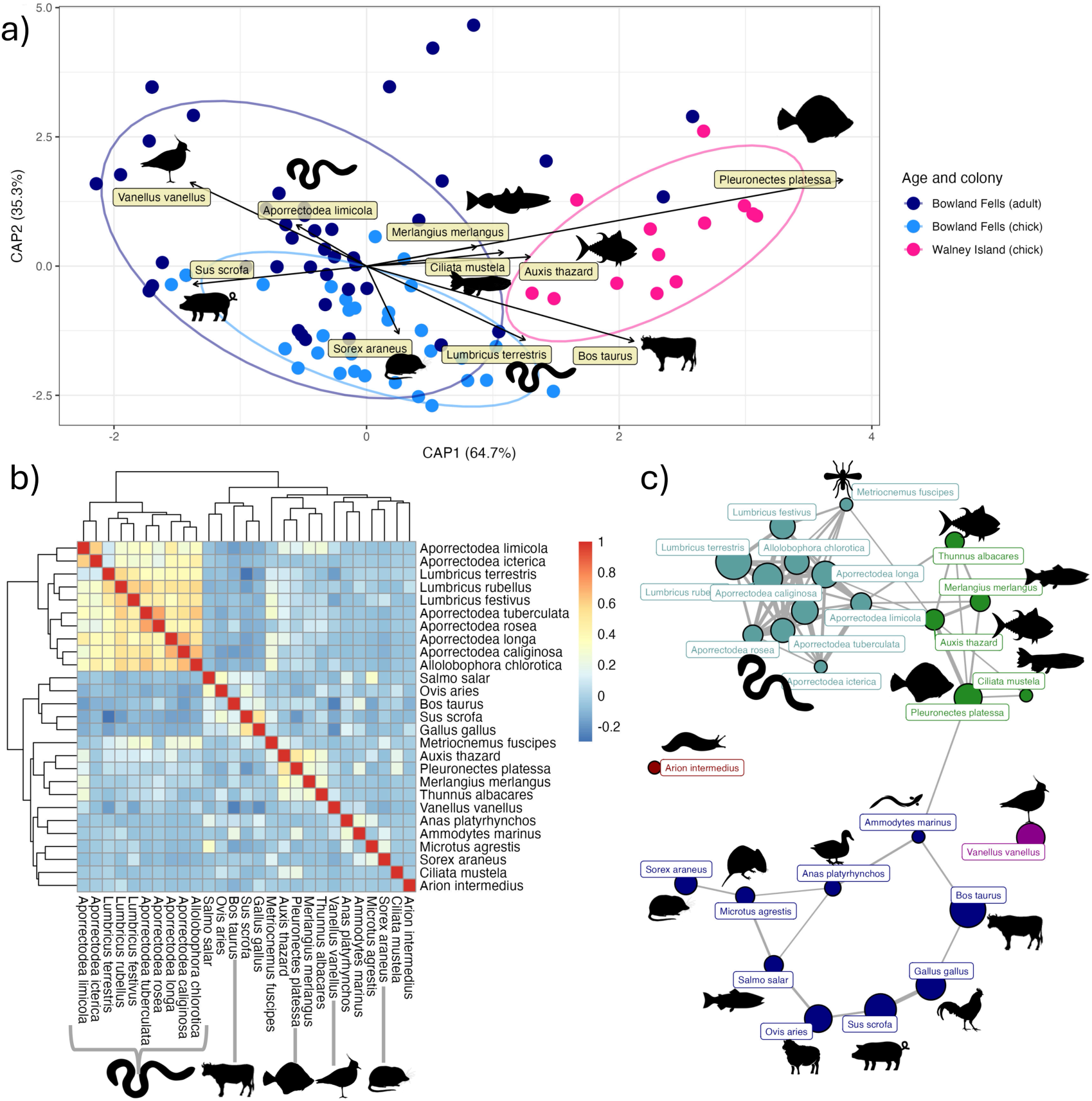
Dietary niche and network analysis. b) A biplot of a constrained ordination visualising compositional differences between age and colony cohorts. Dots represent individual gulls, and arrows represent key species driving differences in dietary niche; b) A heatmap showing co-occurrence of detected species (detected in at least three gulls); c) Network of co-occurring clusters of species shown in (b). Dots represent species and are coloured by cluster membership, and sized by prevalence.

Finally, we conducted a network analysis of prey co-occurrence within individual birds to better understand foraging strategies (Fig. 6b,c). Interpretation of this network may be confounded by the fact that chicks are fed by two parents, each with a potentially contrasting foraging strategy.

The strongest co-occurrence cluster was observed among the 10 common earthworm species (*Lumbricus* and *Aporrectodea* spp.; Fig. 6b & c), representing foraging on agricultural land for earthworms and other invertebrates.

A second cluster included fish species such as European plaice (*P. platessa*), whiting (*M. merlangus*), and genera from the mackerel family (*Thunnus* and *Auxis* spp.). As tuna detections were correlated with those of whiting and plaice, it is possible that some tuna detections may actually represent commercial species such as Atlantic bonito or Atlantic mackerel, as the mackerel family is difficult to distinguish with 12S primers. The association of European plaice with other commercially sourced marine fish species suggests that this flatfish may originate from fishery discards or foraging around fishing vessels rather than direct coastal foraging.

A more complex cluster included species associated with domestic animals, such as cow (*B. taurus;* beef, dairy), sheep (*O. aries;* lamb, mutton), pig (*S. scrofa;* pork), chicken (*G. gallus;* meat and eggs), and salmon (*Salmo salar*), with the strongest link between chicken and pig (Fig. 6, b & c). However, this cluster also included wild prey including the common shrew (*Sorex araneus*) and short-tailed field vole (*M. agrestis*), as well as mallard duck (*Anas platyrhynchos*) and sand eel (*Ammodytes marinus*), which suggest natural foraging strategies, although sand eel can also be found in bird feed. This complex cluster may either be due to small sample sizes, low prevalance of key prey species, and/or mixed foraging strategies of one or both parents. European lapwing, which was detected in some Bowland adults, did not cluster with any other species (Fig. 6c).

## Discussion

Our study evaluated metabarcoding methods to characterise the broad diet of the declining and highly generalist lesser black-backed gull, and examined how metabarcoding data can reveal differences in resource use across age and colony cohorts. Metabarcoding identified a highly generalist diet, detecting 38 vertebrates, 15 arthropods, 13 annelids, and 5 mollusc species, believed to be derived from both natural and anthropogenic sources. Regurgitate samples from chicks showed the highest dietary diversity, while faecal samples yielded similar, though less diverse, data when using one of the 12S blocking primers we developed. Carcass swabs were less effective overall, though stomach-content swabs often enabled species-level identification. Co-occurrence and ordination analyses revealed clusters of co-occurring species presumed to represent distinct foraging strategies, and showed that dietary niche differed across cohorts: coastal (Walney) chicks relied more on fish, especially European plaice, while inland Bowland gulls consumed more human-associated foods, supplemented by free-living small mammals and, in adults, European lapwing. Earthworms were commonly consumed across all cohorts.

### Performance of amplification and blocking primers

Both 12S and COI primers were used to capture vertebrate and non-vertebrate prey. 12S performed well across a broad range of vertebrate taxa, while COI was strongly biased towards invertebrates and detected vertebrates less reliably. Because no blocking primers were developed for COI, it generally performed poorly on faecal samples due to increased host amplification, though it often detected the earthworm *Lumbricus terrestris*, likely reflecting the high prevalence of earthworm DNA across samples, which may have outcompeted vertebrate DNA for amplification. Combining 12S and COI therefore gave the most comprehensive assessment, as each captured complementary dietary information. Additional primer sets could improve resolution further. For instance, fish-specific primers such as miFish (Miya et al., 2015) and plant-targeting primers (Kartzinel et al., 2015) may offer valuable complementary insight, the latter potentially revealing habitat use and anthropogenic resource exploitation via detection of foods like tomato or wheat. Several fish taxa also showed limited taxonomic resolution with the primers used here, particularly with 12S, due to its short amplicon length.

Blocking primers were necessary to characterise diet from faecal samples but were not necessarily required for other sample types, such as regurgitates. We developed and tested three blocking primers, two of which successfully reduced gull DNA amplification. However, the primer with the strongest host suppression also substantially increased amplification of contaminant DNA, potentially producing unreliable data. This is consistent with previous findings that blocking primers can interfere negatively with amplification (Zabala et al., 2025), suggesting that very strong host suppression may increase bias by disproportionately affecting non-target DNA. Blocking primers also performed poorly on intestinal faecal material and cloacal swabs from carcasses, suggesting that variation in DNA degradation across sample types influences their effectiveness, with DNA from carcass samples possibly being too degraded to yield informative data. Although we applied blocking primers across all sample types to standardise methods, we recommend restricting their use to sample types that do not yield usable data without them, given their potential to introduce bias. Where they are used, it is especially important to follow rigorous eDNA-style protocols risk (e.g., processing samples in fume hoods and including many negative controls collected across each step; Dickie et al., 2018) to reduce and account for contamination.

### Performance across sample types

Regurgitate samples yielded a mean of four species per sample (maximum fifteen), compared with a mean of two (maximum six) for faecal samples, using combined 12S and COI data.

This is consistent with faecal samples containing more host DNA, which is accumulated from sloughed intestinal epithelial cells and increased DNA decay during digestion, whereas regurgitates consist largely of undigested food and may better avoid these issues (Thuo et al., 2019). Despite their lower diversity, faecal samples broadly reflected the same population-level dietary patterns as regurgitates, suggesting they do not strongly bias diet composition but instead recover the most abundant dietary DNA while under-detecting rarer prey DNA. This may partly reflect the still-substantial proportion of gull DNA remaining even with the blocking primer used.

Not all study species regurgitate, however, and faecal samples, along with pharyngeal and cloacal/anal swabs, are often the most accessible sample types available. Cloacal swabs from carcasses did not yield informative data, yet pharyngeal swabs successfully detected recently ingested prey still present in the stomach, including European lapwing and short-tailed field vole, suggesting this could be an effective non-lethal method for identifying recent predation. We expect both swab types would perform better on fresh samples from live animals stored appropriately to limit degradation, as demonstrated previously in other dietary specialists, including insectivorous birds (Vo & Jedlicka, 2014), sharks feeding on teleost fish and crustaceans (Ryburn et al., 2026), and turtles eating macrophytes, fish, and invertebrates (Díaz-Abad et al., 2021). Whether generalist predators would still require blocking primers on these sample types remains unclear.

### Identifying key indicator species for distinct foraging strategies across colonies

Lesser black-backed gulls are undergoing complex demographic shifts (Burnell et al., 2023), and understanding resource use during chick-rearing may help identify the drivers of these changes. Walney chicks showed a dietary profile distinct from both chicks and adults at Bowland, with higher frequencies of fish, particularly European plaice. Bowland chicks, in contrast, showed higher frequencies of domestic animal species (e.g. pig, from pork) and small mammals such as common shrew. Both colonies showed high frequencies of earthworms. The recent population increase at Bowland (APEM, 2023; Burnell et al., 2023) may therefore partly reflect reliable earthworm and urban (e.g. landfill) resources nearby, though further sampling at Walney and other colonies would be needed to confirm any link between diet and demographic trends.

We aimed to identify dietary clusters associated with urban, agricultural, and coastal/marine foraging. Co-occurrence analysis revealed candidate markers of urban foraging: chicken, pig, cow, sheep, and salmon detections tended to cluster together, suggesting shared association with anthropogenic resource use. However, the structure of this cluster was complex, indicating not all detections were necessarily driven by urban foraging. Cow detections were less strongly associated with chicken and pig than would be expected if all three primarily derived from urban food waste. Walney chicks, located near cattle pasture, showed high cow DNA but relatively low pig and chicken; Bowland gulls, from an area dominated by sheep grazing, showed higher sheep DNA instead. This raises the possibility that cow and sheep DNA may partly originate from foraging in grazing landscapes (for example via incidental ingestion of livestock dung while feeding on invertebrates) rather than direct consumption of meat or dairy. Overall, chicken and pig appear to be the most reliable indicators of urban or landfill foraging, whereas cow and sheep may, in some cases, instead reflect resource use in habitats with grazing livestock.

We also detected a dietary cluster of fish species, including the European plaice, which was frequently found in samples from Walney chicks. While plaice presence could reflect natural coastal foraging, its clustering with other commercial fish species such as whiting (*Merlangius merlangus*), rather than with coastal prey like crustaceans, suggests it may instead reflect exploitation of discards from commercial fishing vessels in the Irish Sea - a foraging strategy well documented in this species, particularly in western mainland Europe (Tyson et al., 2015). A small number of Walney chicks also showed detections of non-commercial marine or coastal species, including green crab (*Carcinus maenas*), Raitt’s sand eel (*Ammodytes marinus*), five-bearded rockling (*Ciliata mustela*), and greater pipefish (*Syngnathus acus*), although we note that sand eel may originate from bird or livestock feed. These may indicate natural coastal foraging or fishing discards, but were too infrequent to include in the cluster analysis; further sampling at Walney would improve confidence in identifying marine and coastal foraging strategies.

We identified a high diversity of earthworm species linked to agricultural and grassland foraging, with earthworms being an important and well-documented component of gull diet (Coulson & Coulson, 2008). *L. terrestris* and *L. rubellus* were the most common species, found in gulls from both colonies, with Bowland birds (adults and chicks) showing greater diversity than Walney, potentially reflecting more heterogeneous pasture and agricultural land use around the Bowland colony. All identified earthworm species were predominantly associated with farmland habitats, providing strong evidence of agricultural foraging, although *A. limicola* was particularly associated with Bowland. Other invertebrates were detected at much lower frequencies, suggesting more opportunistic or incidental consumption (e.g. via soil ingestion) rather than active targeting. As knowledge of UK earthworm habitat associations and fine-scale distributions continues to improve (Burton et al., 2024), earthworm species composition may become a valuable tool for understanding gull foraging ecology in agricultural landscapes.

Finally, we identified a subset of adult gulls in Bowland (∼18%) that had consumed European lapwing chicks. These birds were taken within wader breeding habitat and as part of wader protection measures, and are therefore unlikely to represent typical foraging strategies of adult gulls at Bowland more broadly. Birds with lapwing remains generally lacked dietary signatures associated with earthworm foraging or urban resource use, and lapwing detections were negatively associated with human-derived food resources, together tentatively suggesting these individuals were actively hunting for breeding birds, rather than opportunistically consuming them while foraging for other prey such as earthworms.

Unbiased sampling of adult gulls to quantify individual foraging strategies would be a valuable area of future research.

### Conclusions

Our study highlights the value of diet metabarcoding for understanding the ecology and impacts of generalist mesopredators, a trophic group central to structuring ecological systems. To fully characterise the broad diet of lesser black-backed gulls and similar generalists, we recommend using multiple primer sets to capture both vertebrate and invertebrate prey alongside anthropogenic resource use. Regurgitate samples performed best at capturing diet diversity, whilst samples from carcasses generally performed poorly, highlighting the need for fresh samples and appropriate sample storage to avoid DNA degradation. Faecal samples required blocking primers, yet we recommend stringent eDNA-level laboratory protocols to manage the contamination and taxonomic bias blocking primers can introduce. Overall, our study provides a framework for the continued application and development of diet metabarcoding approaches to quantify the diet of gulls and other generalist mesopredators, highlighting the urgent need to understand how dietary variation shapes demographic processes in highly dynamic populations.

## Acknowledgements

We would like to sincerely thank all volunteers, landowners, land managers and organisations that facilitated data collection and access, including but not limited to Cumbria Wildlife Trust, Walney Bird Observatory, United Utilities, and RSPB. We would also like to thank Natural England and the Environmental Research and Innovation Centre (University of Salford) for funding.

## SUPPLEMENTARY FIGURES

**Figure S1).**
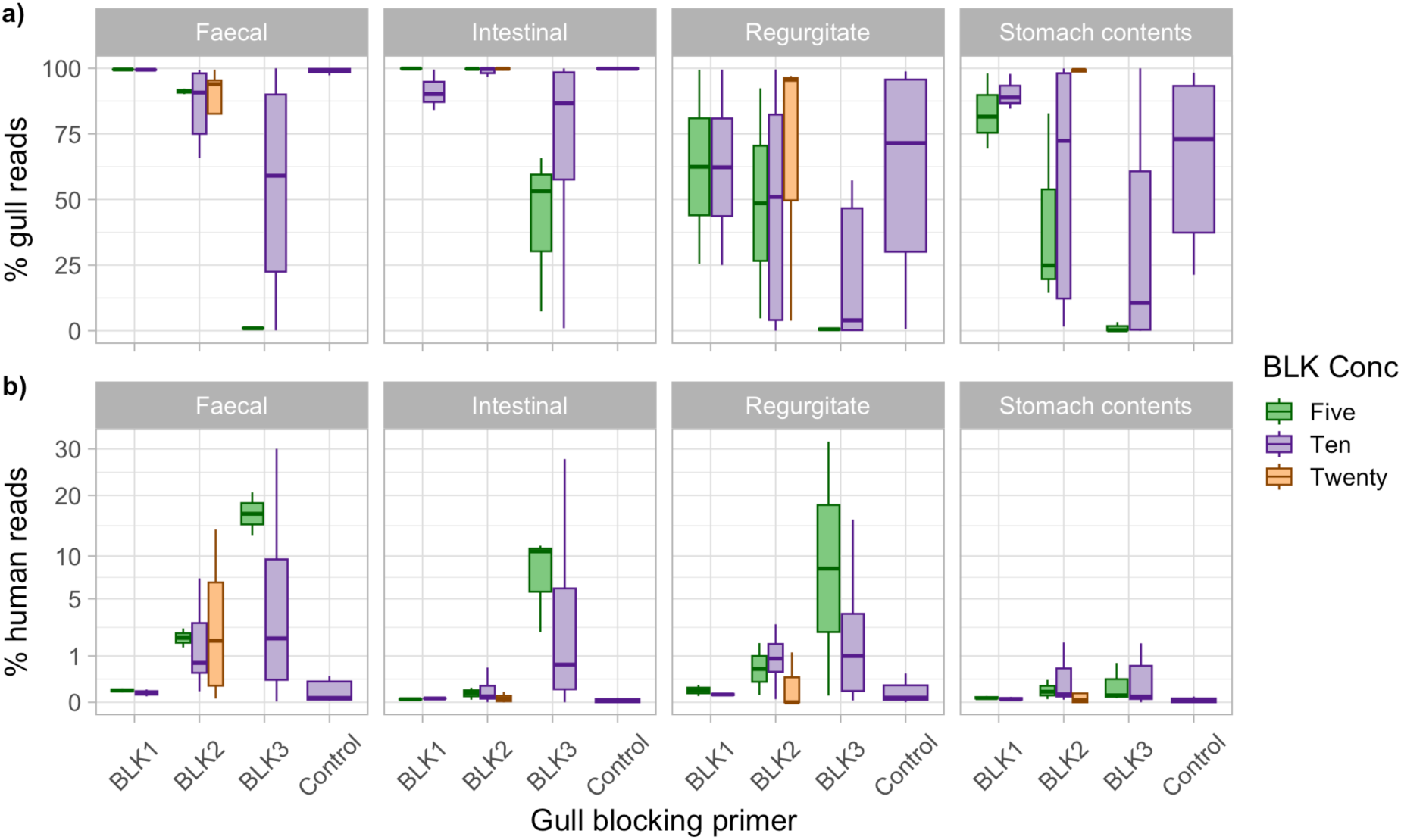
Proportion of gull and human reads across sample types and blocking primers, split by blocking primer concentration. Note not every combination of blocking primer and concentration was tested.

## SUPPLEMENTARY TABLES

**Table S1).**
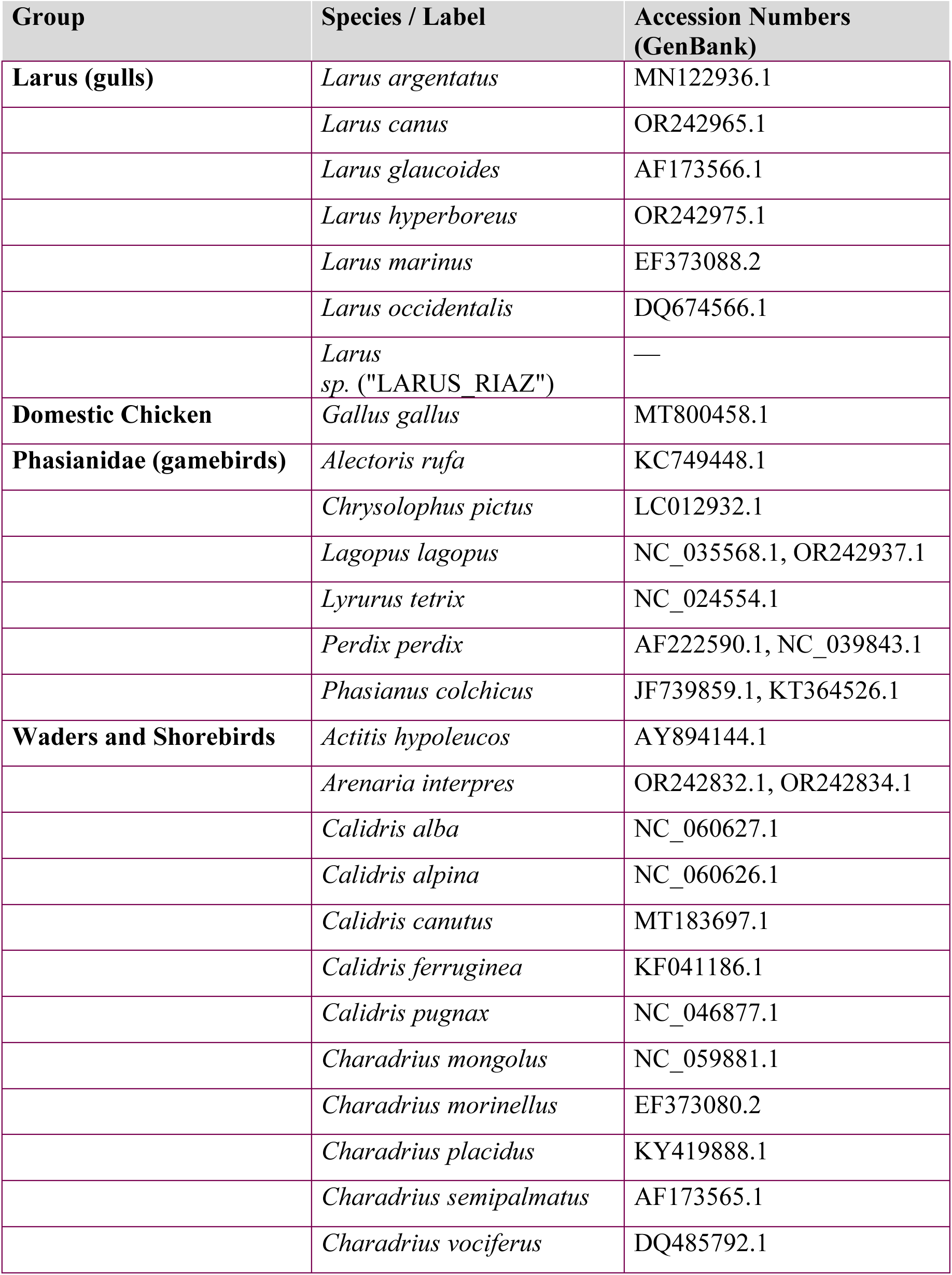
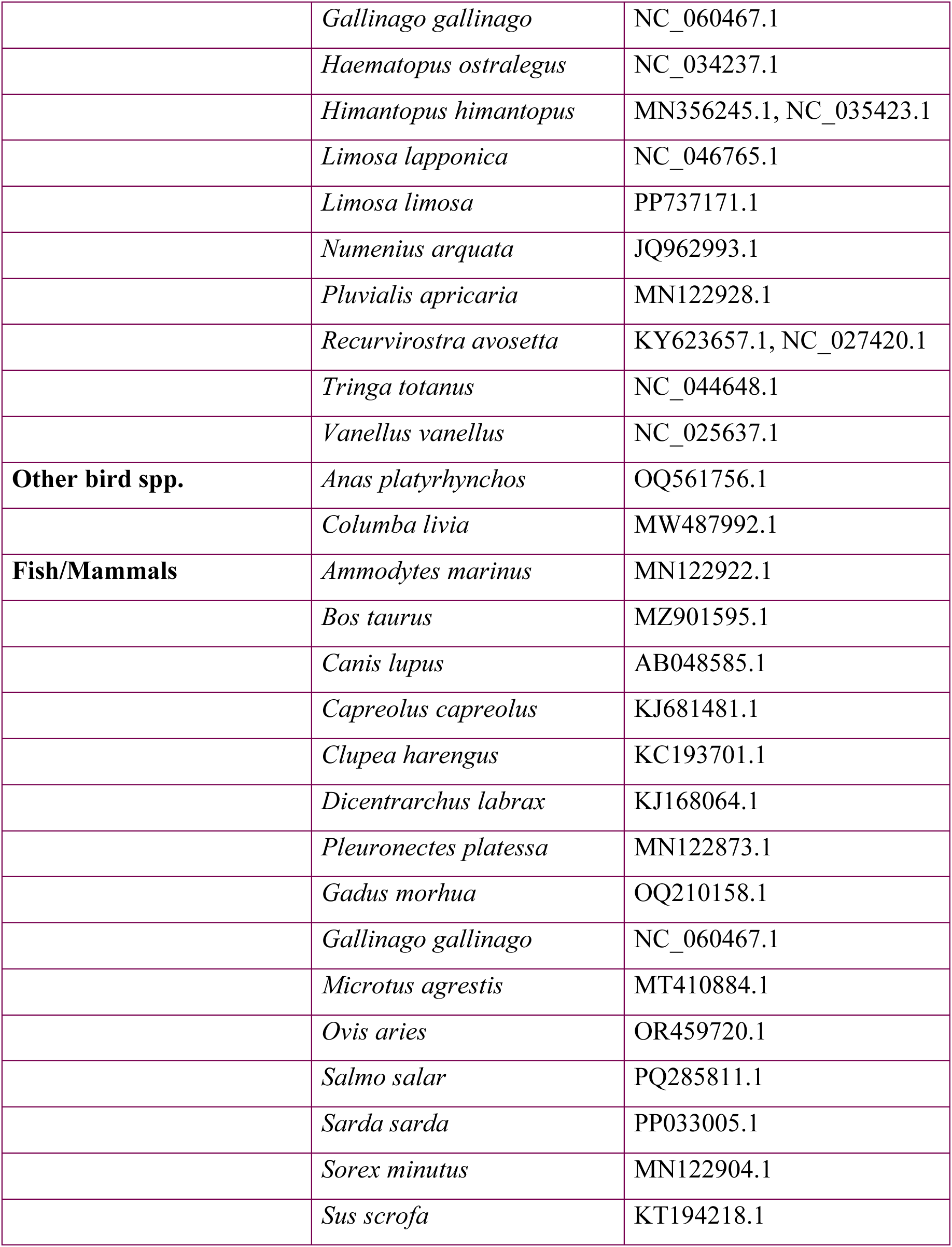
List of species and their corresponding NCBI accession numbers used for the design of blocking primers.

| Group | Species / Label | Accession Numbers (GenBank) |
| --- | --- | --- |
| **Larus (gulls)** | *Larus argentatus* | MN122936.1 |
|  | *Larus canus* | OR242965.1 |
|  | *Larus glaucoides* | AF173566.1 |
|  | *Larus hyperboreus* | OR242975.1 |
|  | *Larus marinus* | EF373088.2 |
|  | *Larus occidentalis* | DQ674566.1 |
|  | *Larus*<br>*sp.* ("LARUS_RIAZ") | — |
| **Domestic Chicken** | *Gallus gallus* | MT800458.1 |
| **Phasianidae (gamebirds)** | *Alectoris rufa* | KC749448.1 |
|  | *Chrysolophus pictus* | LC012932.1 |
|  | *Lagopus lagopus* | NC_035568.1, OR242937.1 |
|  | *Lyrurus tetrix* | NC_024554.1 |
|  | *Perdix perdix* | AF222590.1, NC_039843.1 |
|  | *Phasianus colchicus* | JF739859.1, KT364526.1 |
| **Waders and Shorebirds** | *Actitis hypoleucos* | AY894144.1 |
|  | *Arenaria interpres* | OR242832.1, OR242834.1 |
|  | *Calidris alba* | NC_060627.1 |
|  | *Calidris alpina* | NC_060626.1 |
|  | *Calidris canutus* | MT183697.1 |
|  | *Calidris ferruginea* | KF041186.1 |
|  | *Calidris pugnax* | NC_046877.1 |
|  | *Charadrius mongolus* | NC_059881.1 |
|  | *Charadrius morinellus* | EF373080.2 |
|  | *Charadrius placidus* | KY419888.1 |
|  | *Charadrius semipalmatus* | AF173565.1 |
|  | *Charadrius vociferus* | DQ485792.1 |
|  | *Gallinago gallinago* | NC_060467.1 |
|  | *Haematopus ostralegus* | NC_034237.1 |
|  | *Himantopus himantopus* | MN356245.1, NC_035423.1 |
|  | *Limosa lapponica* | NC_046765.1 |
|  | *Limosa limosa* | PP737171.1 |
|  | *Numenius arquata* | JQ962993.1 |
|  | *Pluvialis apricaria* | MN122928.1 |
|  | *Recurvirostra avosetta* | KY623657.1, NC_027420.1 |
|  | *Tringa totanus* | NC_044648.1 |
|  | *Vanellus vanellus* | NC_025637.1 |
| **Other bird spp.** | *Anas platyrhynchos* | OQ561756.1 |
|  | *Columba livia* | MW487992.1 |
| **Fish/Mammals** | *Ammodytes marinus* | MN122922.1 |
|  | *Bos taurus* | MZ901595.1 |
|  | *Canis lupus* | AB048585.1 |
|  | *Capreolus capreolus* | KJ681481.1 |
|  | *Clupea harengus* | KC193701.1 |
|  | *Dicentrarchus labrax* | KJ168064.1 |
|  | *Pleuronectes platessa* | MN122873.1 |
|  | *Gadus morhua* | OQ210158.1 |
|  | *Gallinago gallinago* | NC_060467.1 |
|  | *Microtus agrestis* | MT410884.1 |
|  | *Ovis aries* | OR459720.1 |
|  | *Salmo salar* | PQ285811.1 |
|  | *Sarda sarda* | PP033005.1 |
|  | *Sorex minutus* | MN122904.1 |
|  | *Sus scrofa* | KT194218.1 |

**Table S2).** General Linear Model outputs for detected species diversity across sample types using a) 12S primers with blocking primer, and b) COI primers.

**a) 12S Diversity**
| *Predictors* | *Incidence Rate Ratios* | *95% CI* | *p* |
| --- | --- | --- | --- |
| (Intercept)[Ref: Pharyngeal] | 0.14 | 0.04 – 0.57 | **0.006** |
| Sample Type [Stomach contents] | 7.66 | 1.89 – 31.08 | **0.004** |
| Sample Type [Intestinal] | 1.6 | 0.38 – 6.77 | 0.523 |
| Sample Type [Regurgitate] | 16.41 | 4.03 – 66.81 | **<0.001** |
| Sample Type [Faecal] | 11.13 | 2.72 – 45.49 | **0.001** |
| Observations (n) | 275 |  |  |
| R<sup>2</sup> | 0.548 |  |  |

**b) COI Diversity**
| *Predictors* | *Incidence Rate Ratios* | *95% CI* | *p* |
| --- | --- | --- | --- |
| (Intercept) | 0.33 | 0.15 – 0.74 | **0.007** |
| Sample Type [Stomach contents] | 5.23 | 2.32 – 11.79 | **<0.001** |
| Sample Type [Intestinal] | 0.28 | 0.10 – 0.77 | **0.014** |
| Sample Type [Regurgitate] | 10.18 | 4.46 – 23.23 | **<0.001** |
| Sample Type [Faecal] | 3 | 1.22 – 7.37 | **0.017** |
| Observations (n) | 268 |  |  |
| R<sup>2</sup> | 0.715 |  |  |

**Table S3).** Results of DNA metabarcoding analysis of samples from dissected gulls in relation to findings from dissections.

| Dissection findings | Total number of birds | Lapwing | Fish | Chicken/<br>pig/<br>cattle/<br>sheep | Earthworms & slugs | Vole or shrew | Hare | No hits |
| --- | --- | --- | --- | --- | --- | --- | --- | --- |
| Lapwing chick remains | 12 | 12 | - | - | 4 | - | - | - |
| Unidentifiable wader chick remains | 4 | 4 | 1 | 1 | - | - | - | - |
| Unidentifiable bird remains | 8 | 2 | 2 | 7 | - | - | - | - |
| Unidentifiable soft material (mush) | 53 | 1 | 3 | 20 | 12 | 1 | - | 20 |
| Other vertebrate | 4 | - | - | 3 | - | 1 | - | - |
| Earthworms | 5 | - | 1 | 2 | 3 | 1 | - | 1 |
| Empty | 21 | - | 1 | 5 | 3 | — | 1 | 14 |
| **Total** | **107** | **19** | **8** | **38** | **22** | **3** | **1** | **35** |
|  |  | **18%** | **7%** | **36%** | **21%** | **3%** | **1%** | **33%** |

